# Inhibition of the Myocardin-Related Transcription Factor pathway increases efficacy of Trametinib in NRAS-mutant melanoma cell lines

**DOI:** 10.1101/773531

**Authors:** Kathryn M. Appleton, Charuta C. Palsuledesai, Sean A. Misek, Maja Blake, Joseph Zagorski, Thomas S. Dexheimer, Richard R. Neubig

**Affiliations:** Department of Pharmacology & Toxicology, Michigan State University, East Lansing, MI; Nicholas V. Perricone, M.D., Division of Dermatology, Department of Medicine, Michigan State University, East Lansing, MI

**Keywords:** NRAS mutant melanoma, drug resistance, Rho/MRTF pathway, MEK inhibitors, MRTF pathway inhibitor

## Abstract

The Ras/MEK/ERK pathway has been the primary focus of targeted therapies in melanoma; it is aberrantly activated in almost 80% of human cutaneous melanomas (∼50% BRAF^V600^ mutations and ∼30% NRAS mutations). While targeted therapies have yielded success in BRAF^V600^ mutant melanoma patients, such therapies have been ineffective in NRAS mutant melanomas in part due to their cytostatic effects and primary resistance in this patient population. Here, we demonstrate that increased Rho/MRTF-pathway activation correlates with high intrinsic resistance to the MEK inhibitor, trametinib, in a panel of NRAS mutant melanoma cell lines. Combination of trametinib with the Rho/MRTF-pathway inhibitor, CCG-222740, synergistically reduced cell viability in NRAS mutant melanoma cell lines *in vitro*. Furthermore, the combination of CCG-222740 with trametinib induced apoptosis and reduced clonogenicity in SK-Mel-147 cells which have a high level of trametinib resistance. These findings suggest a role of the Rho/MRTF-pathway in intrinsic trametinib resistance in a subset of NRAS mutant melanoma cell lines and highlights the potential of concurrently targeting the Rho/MRTF-pathway and MEK in NRAS mutant melanomas.

## INTRODUCTION

With expanding knowledge of the genomic landscape of melanoma, targeted therapies have been evolving over the last decade (1). Almost half of melanoma patients harbor BRAF^V600^ mutations, followed by the second most common oncogenic driver mutations occurring in the NRAS gene in approximately 30% of melanoma patients (2,3). A number of FDA approved targeted therapies have become standard of care in BRAF mutant melanoma patients. Inhibitors targeting mutant BRAF, such as vemurafenib and dabrafenib, were initially employed as monotherapies. In recent years, BRAF/MEK inhibitor combinations including debrafenib/trametinib, vemurafenib/cometinib, and encorafenib/binimetinib have been approved by the FDA, leading to median progression-free survival (PFS) of up to 14.9 months in BRAF mutant melanoma patients (4). Compared to melanoma patients with BRAF^V600^ mutations, patients with activating NRAS mutations are associated with more aggressive disease progression and poor prognosis (5). To date, NRAS mutant melanoma patients have limited treatment options consisting of chemotherapeutics and immunotherapies, both of which are associated with high toxicity (6-9). Unfortunately, targeted therapies for NRAS mutant melanoma patients are sorely lacking.

N-Ras protein targeted small molecule inhibitors have been difficult to design and efforts to inhibit posttranslational modification of N-Ras via farnesyltransferase inhibitors have not yielded approved therapies yet (10-12). Therefore, efforts to develop targeted therapies for NRAS mutant melanomas have focused on signaling components downstream of NRAS in the MAPK pathway, such as mitogen-activated protein kinase kinase (MEK). Only a subset of patients benefits from MEK inhibitors used as single agents, because they only produce cytostatic effects rather than cytotoxic effects, in NRAS mutant melanoma cells (13). MEK inhibitors are also associated with primary and acquired resistance as well as frequent toxicity-related adverse events (14,15). However, MEK inhibitors could deliver promising therapies when combined with inhibitors of other signaling mechanisms (12,15-17). Currently, the MEK inhibitor FCN-159, which has 10-fold higher selectivity against activated MEK1/2 compared to trametinib, is being investigated as a single agent in a phase I clinical trial (NCT03932253). Additionally, clinical trials for the MEK inhibitor trametinib in combination with an ErbB3 antibody (NCT03580382) or a pan-RAF inhibitor, LXH254, (NCT02974725) are currently underway. However, all of these studies are still in early-phase clinical trials. Furthermore, primary and acquired resistance to some of the combinations, such as the MEK inhibitor MEK162 with a CDK4/6 inhibitor (18,19), are emerging. Therefore, there is a continued need to understand the mechanism of resistance of NRAS mutant melanomas to MEK inhibitors and to develop new treatment strategies.

In cutaneous melanoma patients, increased expression of RhoC or MRTF-A mRNA has been linked to overall poor patient survival (20). Myocardin-related transcription factor (MRTF) is a pair (MRTF-A and -B) of transcription cofactors downstream of Rho GTPases (21,22). Rho GTPases regulate the actin cytoskeleton and have long been demonstrated to play a critical role in cellular invasion and metastasis in numerous human cancers including melanoma (23,24). Activation of RhoA and RhoC GTPases causes subsequent activation of their effector protein, Rho-associated protein kinase (ROCK), which leads to formation of F-actin polymers and resultant depletion of free G-actin monomers (25). In the cytosol, G-actin monomer binds to the RPEL domain in the N-terminal region of MRTF and sequesters MRTF away from the nucleus (26). Upon F-actin polymerization during the formation of stress fibers, MRTF translocates to the nucleus, where it cooperates with serum response factor (SRF) to induce transcription of numerous genes involved in cell proliferation and migration (21,22). The MRTF-SRF transcriptional axis plays a pro-metastatic role in the context of melanoma and other cancers (27). Depletion of MRTF via RNA interference (RNAi) in the highly metastatic B16F2 melanoma cell line reduced *in vitro* cell migration and *in vivo* lung metastasis (26). Additionally, pharmacologic inhibition of the Rho/MRTF-pathway by the small-molecule CCG-203971 significantly reduced *in vitro* cellular migration and invasion (20,28), as well as *in vivo* lung metastasis in the RhoC-expressing NRAS mutant melanoma cell line SK-Mel-147 (20). In addition to its anti-migratory and anti-metastatic properties, CCG-203971 induced G1-cell cycle arrest in melanoma cells. The recent observation that melanoma cells arrested in the G1 phase have higher sensitivity to MEK inhibitors (29), suggests a potential benefit of combination treatment with MEK inhibitors and Rho/MRTF pathway inhibitors.

Through structure-activity relationship optimization of CCG-203971, we recently reported the analog CCG-222740 with increased potency of MRTF-pathway inhibition in primary human dermal fibroblasts (30). Additionally, CCG-222740 demonstrated a greater inhibitory effect on MRTF/SRF target genes (ACTA2 and CTGF) and lesser cytotoxicity than CCG-203971 in a preclinical model of fibrosis (31). Building on our previous work showing that CCG-203971 inhibited melanoma metastasis, here, we evaluated the pharmacological potential of CCG-222740, a more potent CCG-203971 analog, in combination with a MEK inhibitor (trametinib) in NRAS-mutant melanoma cells. In a panel of NRAS mutant melanoma cell lines, we observed a correlation between the degree of activation of the Rho/MRTF pathway and intrinsic resistance of cells to trametinib-mediated apoptosis. CCG-222740 potentiates trametinib action in the subset of NRAS mutant melanoma cells which had a high level of activation of the Rho/MRTF pathway. In these cells, the combination of trametinib and CCG-222740 cooperatively induced apoptosis and reduced colony formation potential.

## MATERIALS AND METHODS

### Inhibitors

CCG-222740, previously reported by Hutchings et al. (30), was a kind gift of Scott Larsen at the Vahltechich Medicinal Chemistry Core (Ann Arbor, MI). CCG-222740 was dissolved in DMSO in 10 mM aliquots and stored at −20 °C. Trametinib (cat# S2673), Vinblastine (cat# S1248), Y-27632 (cat# S1049) were purchased from Selleck Chemicals (Houston, TX), reconstituted in DMSO, and 10 mM aliquots were stored at −20 °C.

### Cell Culture and Viability Assay

Human cutaneous melanoma cell lines WM-3451 and WM-3623 were purchased from The Wistar Institute (Philadelphia, PA), SK-Mel-2 cells were purchased from ATCC (Manassas, VA) and SK-Mel-19 and SK-Mel-147 were obtained from Dr. Maria Soengas at the University of Michigan and have been described previously (20). Cells were cultured in DMEM (Life Technologies) supplemented with 10% fetal bovine serum and 1X antibiotic-antimycotic solution (Life Technologies). Cells were expanded and frozen immediately prior to authentication and then thawed only two to three months before experiments. Short tandem repeat profiles were performed on SK-Mel-19 and SK-Mel-147 cell lines (Genewiz, South Plainfield, NJ). The profiles obtained do not match any established published profiles, and we were unable to identify published profiles for SK-Mel-19 or SK-Mel-147. NRAS exon 3 and BRAF exon 15 were PCR amplified from genomic DNA and subjected Sanger sequencing (MSU Genomics Core). The mutation status of the cells used were: NRAS^Q61L^ (SK-Mel-2, SK-Mel-147, WM-3451), NRAS^Q61K^ (WM-3623), BRAF^V600E^ (SK-Mel-19). For viability assays, 1,000 cells in 20 µL of DMEM containing 10% FBS were seeded into 348-well white bottom plates. Four to six hours later, 10 µL of 4X compound was added along with an additional 10 µL of either 4X second compound or media. After 72 hours, 20 µL of CellTiterGlo (Promega) was added to each well. The assay plate was centrifuged at 300 x g for 3 minutes. Luminescence was measured using a BioTek Synergy Neo plate reader. Data were normalized to values obtained for the vehicle-treated cells. Data were plotted as average values of at least three independent experiments. Non-linear least squares analysis was used to fit data to a 4-parameter log-[inhibitor] vs. response curve using GraphPad Prism versions 6-8 (GraphPad Software, La Jolla, CA, USA).

### Clonogenicity Assay

Two hundred cells in DMEM containing 10% FBS were seeded in 6-well dishes and simultaneously treated with vehicle or 6 µM CCG-222740 with or without 0.1 nM or 1 nM Trametinib or with Trametinib alone. Following five days of colony formation, fresh media and compound were added, and colonies were allowed to grow for an additional five days. Colonies were fixed and stained in 3.7% formaldehyde/0.5% crystal violet for 10 minutes at room temperature. Colony counts were quantified using ImageJ software with a cutoff for colony size (pixel^2) fixed at 50-infinity and circularity defined as 0.2-1.0.

### Immunofluorescence and F-actin staining

Cells (5×10^4^) were seeded into Falcon 8 Chamber Slides in 500 µL of DMEM containing 10% FBS and allowed to adhere overnight. Cells were fixed with 3.7% formaldehyde for 10 minutes at room temperature. Chambers were washed 3 times with 1X PBS for 5 minutes each. Cells were permeabilized with 0.25% Triton X-100 in PBS for 10 minutes followed by 3 5-minute PBS washes. For stress fiber detection, rhodamine phalloidin (Cytoskeleton, Inc. cat# PHDR1) was used according to manufacturer’s recommendations. Stress Fibers were scored with slight modification from the reference Verderame et al. (32). Score values followed the criteria: 5: >90% of cell area filled with thick cables; 4: At least two thick cables running under the nucleus and rest of area filled with fine cables; 3: No thick cables, but some fine cables present; 2: No cables visible in center area of cell; 1: No cable visible.

For MRTF-A staining, 10% donkey serum in PBS was used as a blocking solution for 30 minutes. For MRTF-B and YAP staining, 5% bovine serum albumin (BSA) in PBS was used as a blocking solution for 30 minutes. Primary antibodies for MRTF-A (Santa Cruz, cat# sc-21558), MRTF-B (Santa Cruz, cat# sc-98989), or YAP (Santa Cruz cat# sc-15407) were diluted 1:100 in 1% BSA, 0.01% Triton X-100 in PBS and incubated at 4 °C overnight. Following 3 5-minute PBS washes, secondary antibodies for MRTF-A (Alexa-Fluor® 594 donkey anti-goat IgG antibody, Life Technologies cat# A11058), for MRTF-B or YAP (Alexa-Fluor® 594 goat anti-rabbit IgG antibody, Life Technologies cat#A11037) were diluted 1:1,000 in same primary antibody buffer and incubated 1 hour at room temperature. Following 3 5-minute PBS washes, cells were mounted (Prolong Gold antifade-reagent with DAPI, Invitrogen, cat# P36935) and imaged on an EVOS FL Cell Imaging System (Life Technologies) at 40X magnification. All scoring of stress fibers and nuclear localization of transcription factors was done by an observer who was blinded to the identity of the sample.

### Real-time quantitative PCR (qRT-PCR)

Cells (2×10^5^) were seeded into 6-well plates and left to adhere overnight then harvested the following day for CYR61 mRNA analysis. Briefly, RNA was isolated using the RNeasy kit (Qiagen) following manufacturer’s directions. RNA (1 µg) was used as a template for synthesizing cDNA utilizing Reverse Transcriptase Kit (Life Technologies cat#4368814). Real-time PCR was conducted using SYBR Green Master Mix (Life Technologies cat#4309155) using 4 µL of cDNA reaction on a Stratagene Mx3000P instrument (Agilent Technologies, Santa Clara, CA) and cT values were analyzed relative to GAPDH expression. SK-Mel-19 cells, which have a low degree of Rho activation (20), were used to normalize relative CYR61 mRNA levels for testing NRAS mutated cell lines. For SRF mRNA experiments, cells were treated with 2 nM trametinib and harvested at specified time points. The DMSO control was used to normalize relative expression. To assess the effects of CCG-222740 on CYR61, CRIM1, THBS1 and ANKRD1 mRNA levels, 4.5×10^5^ cells were seeded into 6-well plates and left to adhere overnight. Cells were then treated with 10 µM CCG-222740 for 24 hours. RNA isolation and cDNA synthesis were performed as mentioned above. Real-time PCR was conducted using SYBR Green Master Mix and 2 µL of cDNA on the Stratagene Mx3000P instrument. cT values were normalized relative to GAPDH expression, and fold-change in mRNA expression upon CCG-222740 treatment was calculated relative to vehicle control (0.02% DMSO). Primer Sequences: CYR61 F1: 5’-GAAGGTGAAGGTCGGAGTCA-3’; CYR61 R1: 5’-TTGAGGTCAATGAAGGGGTC-3’; CYR61 F2: 5’-CTCGCCTTAGTCGTCACCC-3’; CYR61 R2: 5’-CGCCGAAGTTGCATTCCAG-3’; GAPDH F1: 5’-CTCGCGGCTTACCGACTG-3’; GAPDH R1: 5’-GGCTCTGCTTCTCTAGCCTG-3’; GAPDH F2: 5’-GGAGCGAGATCCCTCCAAAAT-3’; GAPDH R2: 5’-GGCTGTTGTCATACTTCTCATGG-3’; SRF F: 5’-CCTTCAGCAAGAGGAAGACG-3’; SRF R: 5’-GATCATGGGCTGCAGTTTTC-3’; CRIM1 F: GGTTCCTGTTGTGCTCTTGT; CRIM1 R: TGCCAAGAATCAAGTTGCAGATAA; THBS1 F: AGACTCCGCATCGCAAAGG; THBS1 R: TCACCACGTTGTTGTCAAGGG; ANKRD1 F: AGAACTGTGCTGGGAAGACG; ANKRD1 R: GCCATGCCTTCAAAATGCCA.

### Western Blot Analysis

Cells (8×10^5^) were seeded into 60-mm dishes. For both pERK1/2 and pMLC (Phospho-Myosin Light Chain 2, Thr18/Ser19) immunoblots, cells were harvested 24 hours after seeding by direct lysis in 2X Laemmli buffer mixed with equal volume of RIPA buffer (10 mM Tris-Cl pH 8.0, 1mM EDTA, 1% Triton X-100, 0.1% sodium deoxycholate, 0.1% SDS, 140 mM NaCl, protease/phosphatase inhibitor cocktail Thermo Fisher cat#88266). Samples were sonicated 2x for 5 seconds and heated for 5 minutes at 95°C. Equal volumes of protein lysate (30 µL) for each cell line were resolved using 15% SDS-PAGE gels, transferred to PVDF membranes and blocked in 5% BSA in Tween tris-buffered saline. Primary and secondary antibodies were diluted in 1% BSA, 0.01% Triton X-100 in PBS. Membranes were incubated in 1:1000 diluted primary antibodies pERK1/2 pT202/Y204 (Calbiochem cat# KP26001), total ERK1/2 (Cell Signaling cat# 9102), pMLC (Cell Signaling cat# 3674S), GAPDH (Santa Cruz cat#365062) overnight at 4 °C. HRP-conjugated anti-rabbit (Cell Signaling cat# 7074P2), and HRP-conjugated anti-mouse (Cell Signaling cat# 7076P2) antibodies were diluted 1:6000. Blots were developed using FemtoGlow™ Western chemiluminescent HRP substrate (Michigan Diagnostics cat# FWPS02). Bands were visualized and quantified using a Li-Cor Odyssey Fc scanner and quantified using Image Studio Lite Version 5.2.

### Caspase 3/7 Activity Assay

Cells (2×10^5^) were plated in 6-well plates in DMEM with 10% FBS and allowed to adhere overnight. Treatments were done the next day with either 10 µM CCG-222740 and 12.5 nM Trametinib alone or in combination and 0.2% DMSO was used as a vehicle control. CellEvent™ Caspase-3/7 Green Flow Cytometry Assay Kit (Invitrogen) was used according to the manufacturer’s protocol with modest modifications. Briefly, media and cells were harvested using a cell scraper 48 hours after treatment started and centrifuged 300xg for 5 minutes. Cell pellets were washed with PBS and pelleted again. Cells were resuspended in 1 mL of PBS and 150 µL of the cell solution was added to a V-bottom 96-well plate. The plate was centrifuged at 300 x g for 5 minutes and the PBS supernatant was removed. Single-stained compensation controls were included for each cell line for every experiment. 25 µL of either FACs buffer (1% FBS in PBS), or CellEvent™ Caspase-3/7 Green Detection Reagent (diluted 1:1000 in FACS buffer) was added to appropriate wells, the plate was covered with film, and incubated at 37 °C at 5% CO_2_ for 25 minutes. The plate was removed, and 25 µL of either FACs buffer or SYTOX™ AADvanced™ dead cell stain (diluted 1:1000 in FACS buffer) was added to appropriate wells. The plate was covered with film and incubated at 37 °C for 5 minutes. Following incubation, 100 µL of FACs buffer was added to each well and samples were transferred to 0.5 mL Eppendorf tubes in a 5% CO_2_ atmosphere. Fluorescence was detected using a C6 BD Accuri flow cytometer (BD Accuri, San Jose, CA). Fluorescence was quantified using CFLow software (BD Accuri). Events (20,000) were detected in triplicate for all treatment groups and three independent experiments were conducted, and results were averaged.

### RNA-Seq sample preparation and data processing

SK-Mel-147 cells were treated -/+ 10 µM CCG-222740 or vehicle control for 24 h. Total cellular RNA was extracted from (two biological replicates per treatment condition) using the Qiagen, Hilden, Germany, RNeasy kit (#74104) according to the manufacturer’s protocol. RNA was eluted in nuclease-free H_2_O. RNA concentration was measured by Qubit and quality control was performed on an Agilent 2100 Bioanalyzer in the MSU Genomics Core. All RNA samples had a RIN score > 8. Barcoded libraries were prepared using the Illumina TruSeq Stranded mRNA Library Preparation Kit on a Perkin Elmer Sciclone G3 robot following manufacturer’s recommendations. Completed libraries were QC’d and quantified using a combination of Qubit dsDNA HS and Caliper LabChipGX HS DNA assays. Libraries were pooled and run on two lanes, and sequencing was performed in a 1×50 bp single-end read format using HiSeq 4000 SBS reagents. Base calling was done by Illumina Real Time Analysis, RTA_ v2.7.7 and output of RTA was demultiplexed and converted to FastQ format with Illumina Bcl2fastq v2.19.0. Sequencing was performed at a depth of >30M reads/sample. Quality control was performed on the FastQ files using FastQC v0.11.5, and reads were trimmed using Trimmomatic v0.33. Reads were mapped using HISAT2 v2.1.0 and analyzed using HTSeq v0.6.1. Differential gene expression was calculated using edgeR. Raw RNA-Seq reads and processed HTSeq read counts are available on GEO under GSE134320.

## RESULTS

### MRTF pathway activation correlates with trametinib resistance

Trametinib is reported to induce apoptosis at varying levels and is overall less potent in NRAS mutant melanoma cells compared to BRAF mutant melanoma cells (5). We therefore first determined the sensitivity of a panel of four NRAS mutant melanoma cell lines to inhibition of cell viability by trametinib treatment. In addition, we compared them to a BRAF^V600E^ mutant melanoma cell line, SK-Mel-19. The cell line panel included SK-Mel-147, SK-Mel-2, and WM-3451 cells containing NRAS^Q61L^ mutations, and WM-3451 cells containing an NRAS^Q61K^ mutation. All NRAS mutant melanoma cell lines in our panel were less sensitive to trametinib treatment than were the BRAF mutant SK-Mel-19 cells (Fig. 1A). The area-under-the-curve (AUC) determined from the log[trametinib] concentration response curves (−11 to −7) demonstrated significantly greater trametinib resistance as well as significant variability among NRAS mutant cells compared to SK-Mel-19 (Fig. 1B). Melanoma cells harboring NRAS^Q61^ mutations have previously been shown to have hyperactivation of the MAPK pathway (33). Therefore, we quantified pERK1/2 via western blot under basal conditions for each cell line (Fig. 1C). We found that the levels of pERK1/2 varied substantially across the cell line panel with decreasing amounts of pERK1/2 associated with increased trametinib resistance (Fig. 1D).

**Figure 1.**
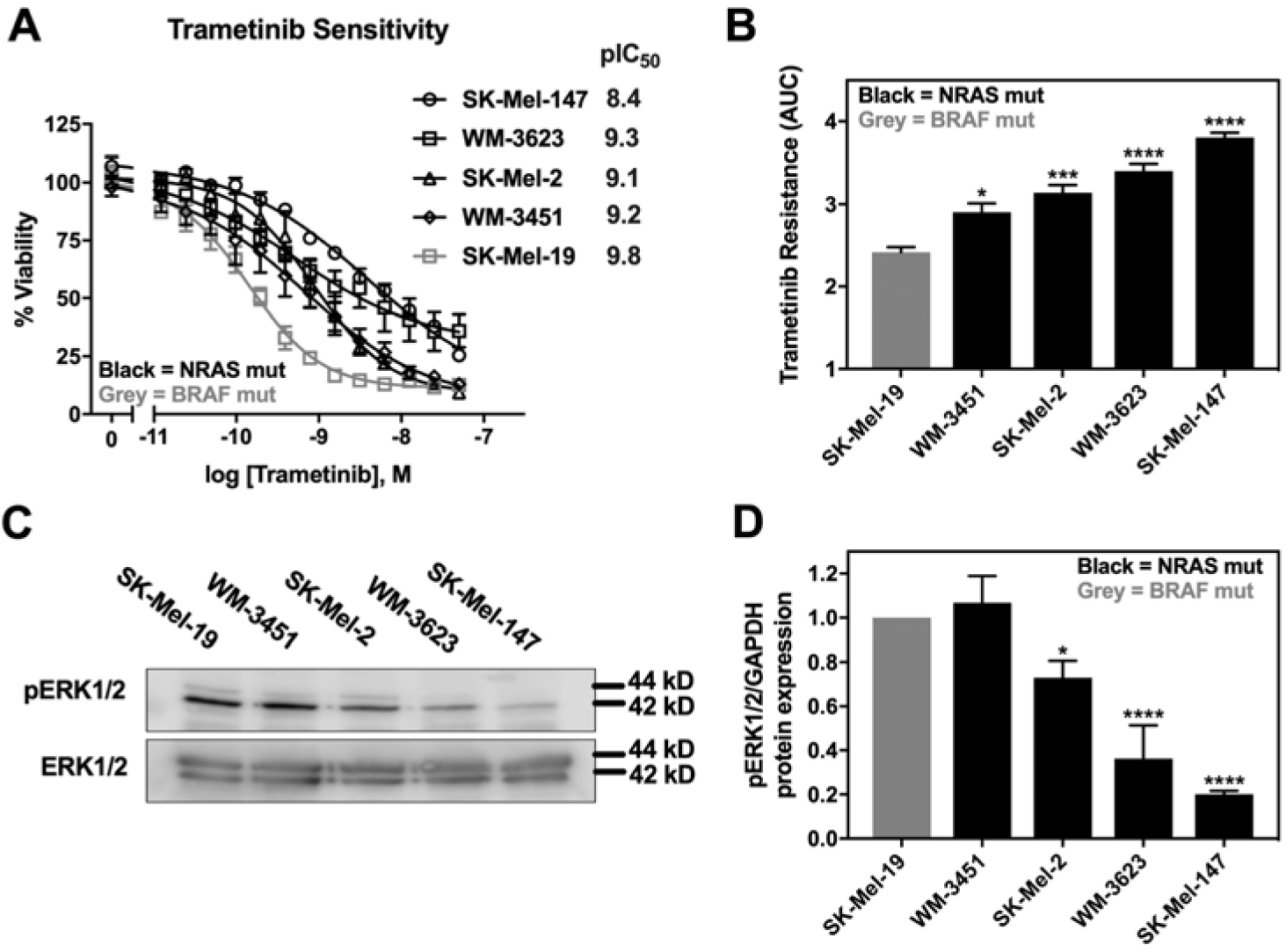
NRAS mutant melanoma cell lines have varying levels of intrinsic resistance to trametinib-induced cell death. **A.** Sensitivity to trametinib was determined by treating each cell line in 10% FBS with increasing concentrations of trametinib for 72 hours. Cell viability was determined using Cell-TiterGLO®. Values in the graph are expressed as the fraction of luminescence vs. vehicle control for at least three independent experiments. pIC_50_ values (-log IC_50_, M) values are indicated for each cell line. **B.** The area under the curve (AUC) was plotted based on the log concentration response curves (log[Trametinib] −11 to −7) generated in panel A using GraphPad Prism. A greater area means less response to trametinib or increased trametinib resistance (*p<0.05, ***p<0.01, ****p<0.001 vs. SK-Mel-19; *p<0.05, WM-3451 vs. SK-Mel-19; **p<0.01, WM-3623 vs. WM-3451; *p<0.05, SK-Mel-147 vs. WM-3623). **C.** Western blot analysis of pERK1/2 across the cell line panel. Each melanoma cell line was plated in 60mm dishes in 10% FBS and harvested 24 hours later.The image is a representative blot from three independent experiments. **D.** Quantitative band density analysis was performed for each experiment comparing the intensity of pERK1/2 relative to total ERK1/2. Results are expressed as the mean (±SEM) of triplicate experiments (*p<0.05, ****p<0.001 vs. SK-Mel-19).

Our recent findings indicated a role of the Rho/MRTF pathway in migration, invasion and metastasis of aggressive human cutaneous melanoma (20,34), as well as in the acquired resistance of BRAF mutant melanoma cells to BRAF inhibitors (34). Therefore, we hypothesized that the Rho/MRTF pathway might play a role in the intrinsic resistance of some of the NRAS mutant melanoma cells to trametinib-induced inhibition of cell viability. Upon activation of the Rho/MRTF pathway, myosin light chain (MLC) gets phosphorylated and G-actin polymerizes into F-actin stress fibers (35). Depletion of G-actin during stress fiber formation results in MRTF translocation into the nucleus where it regulates gene transcription, including an increase in cysteine-rich angiogenic inducer 61 (CYR61) (36). We observed elevated levels of phosphorylated MLC (pMLC) in NRAS mutant cell lines compared to SK-Mel-19, with statistically significantly higher levels of pMLC in SK-Mel-2 cells (Supp. Fig.1). We then stained the cells for actin and scored the images for stress fiber positive cells as described in Methods. Similar to pERK1/2, we detected increased stress fiber formation in NRAS mutant cell lines compared to the BRAF mutant cell line SK-Mel-19. Among NRAS mutant cell lines, SK-Mel-147 cells showed the strongest stress fiber positivity (Fig. 2A-B). When we compared the percentage of stress fiber positive cells to trametinib resistance (AUC), we detected a strong positive correlation (r^2^=0.96, p=0.003) (Fig. 2C). This correlation was still strong when just comparing among NRAS mutant cell lines (r^2^=0.95, p=0.024) Similarly, a significant correlation between the levels of CYR61 mRNA levels and resistance to trametinib (AUC) was also observed for all cells (r2=0.92, p=0.012) and for just NRAS mutant cells (r2=0.74, p=0.0003) (Figure 2D). Since CYR61 gene expression could be regulated upon activation of either the Rho/MRTF pathway via nuclear MRTF-A/B or the Hippo pathway via nuclear YAP, respectively (37), we next studied localization of these transcription factors. Either MRTF-A or MRTF-B were strongly nuclear in 3 of the 4 NRAS mutant cell lines, but not in WM-3451 or SK-Mel-19 cells (Fig. 2E-F, Supp. Fig. 2A). Interestingly, WM-3623 only had MRTF-B in the nucleus but not MRTF-A (see Discussion). Finally, we detected a high percentage of cells with nuclear YAP in our cell panel, including the BRAF mutant SK-Mel-19 cells, with no significant differences across the cell lines (Fig. 2G and Supp. Fig. 2B). Taken together, these results indicate that Rho/MRTF pathway is activated in NRAS mutant melanoma cell lines having intrinsic resistance to trametinib-induced inhibition of cell proliferation.

**Figure 2.**
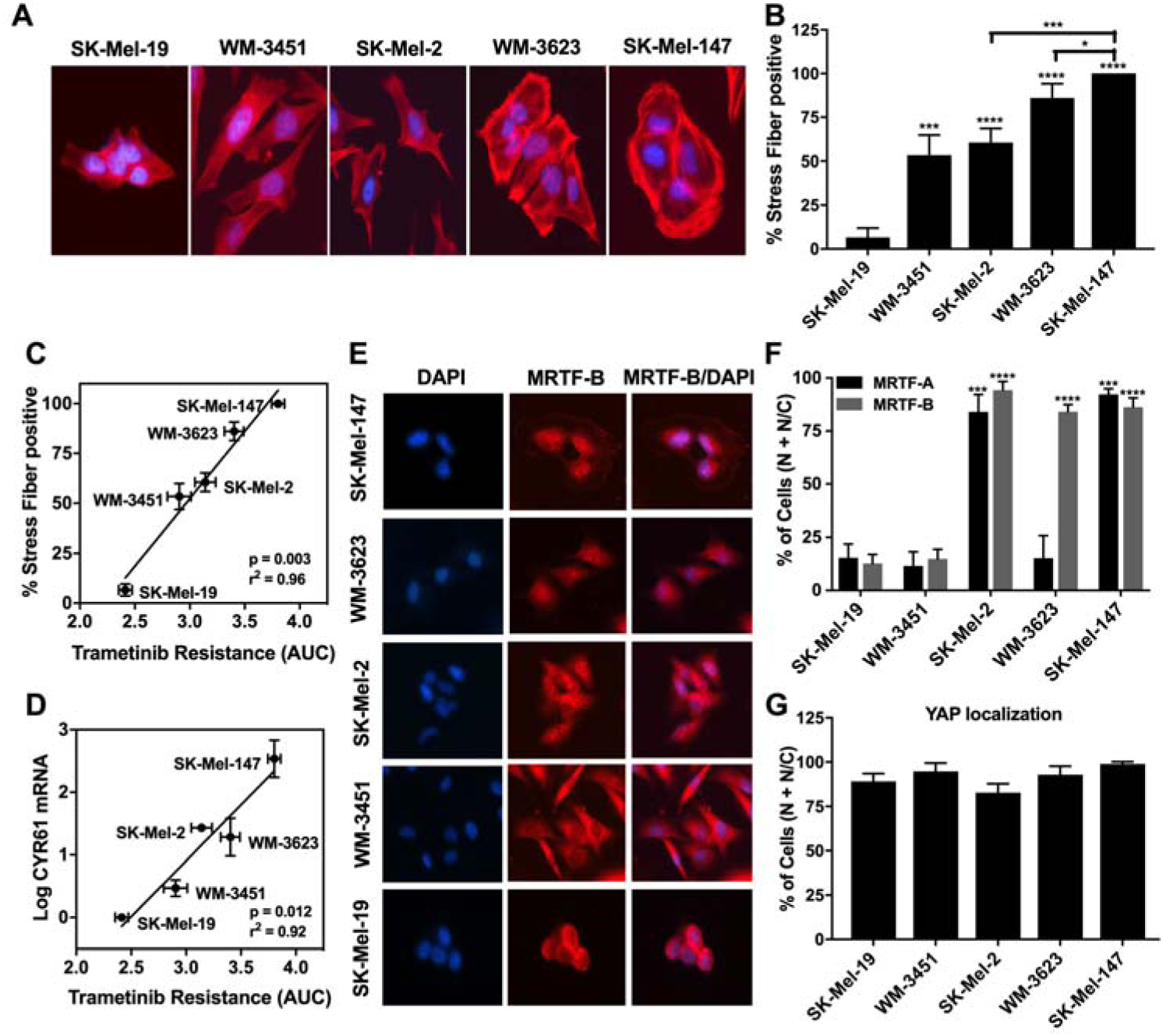
MRTF pathway activation correlates with trametinib resistance. **A.** Actin staining (red) wa assessed on the melanoma cell line panel using fluorophore-conjugated phalloidin toxin. **B.** For stres fiber quantification, a score between 1 to 5 as described in methods was utilized and the percentage of cells scored 3-5 is presented as stress fiber positive for each cell line. Results are expressed as the mean (SEM) of triplicate experiments (*p<0.05, ***p<0.005, ****p<0.001 vs. SK-Mel-19 or SK-Mel-147). **C.** Trametinib resistance correlates with % stress fiber positivity. Utilizing the values depicted in Figure 1B and Figure 2B, a strong positive correlation was observed (r^2^= 0.96, p=0.003, Pearson correlation coefficients, GraphPad Prism). **D.** CYR61 mRNA levels correlates with Trametinib resistance. Utilizing the values depicted in Figure 2D compared to the normalized log CYR61 mRNA expression levels determined using qPCR, a significant positive correlation was observed (r^2^= 0.71, p=0.04, Pearson correlation coefficients, GraphPad Prism). **E.** Representative images of the cellular localization of MRTF-B using immunofluorescence in melanoma cells (10% FBS). **F.** Cellular localization of MRTF-A or MRTF-B observed with immunocytochemistry of melanoma cells. Images were quantified by scoring individual cells as exclusively nuclear (N), cytosolic (C), or even distribution (N/C) of MRTF-A (black bars) or MRTF-B (grey bars). Counts from three independent experiments with at least 100 cells scored for each cell line (***p<0.005, ****p<0.001 vs. SK-Mel-19). **G.** Cellular localization of YAP using immunofluorescence wa determined as stated above for MRTF isoforms. No significant difference was observed across the melanoma cell lines.

### MRTF pathway inhibitor, CCG-222740, synergizes with trametinib to reduce viability of NRAS mutant melanoma cells

Considering a potential role of Rho/MRTF pathway activation in the intrinsic resistance of NRAS mutant cell lines to trametinib treatment, we next sought to determine if the Rho/MRTF-pathway inhibitor CCG-222740 could potentiate the effects of trametinib on viability in our NRAS mutant cell line panel (Fig. 3A). Using cross-concentration response experiments, we observed that SK-Mel-147 cells demonstrated a significant leftward shift in log IC_50_, with a nearly 2 log-shift at the highest concentration of CCG-222740 utilized (Fig. 3A-B). WM-3623 cells and SK-Mel-2 cells also displayed potentiation of trametinib effects when trametinib was combined with CCG-222740 (see Δlog IC_50_ plots, Fig. 3A-B). WM-3451 cells, which had minimal evidence for Rho/MRTF activation, did not display potentiation of trametinib when CCG-222740 was used in combination (Fig. 3A-B). To determine if the observed leftward shifts in log IC_50_ of trametinib in combination with CCG-222740 represents synergistic effects, we calculated the Loewe Excess in order to derive a synergy score (38). Synergy (Loewe Excess 1.67-3.03) was observed for the three NRAS mutant cell lines with the greatest intrinsic trametinib resistance, as well as the greatest detected Rho/MRTF-pathway activity. On the other hand, WM-3451 cells did not display evidence of synergy (Fig. 3C). These results demonstrate that trametinib and CCG-222740 cooperatively reduced the viability of the subset of NRAS mutant melanoma cells lines which have increased Rho/MRTF-pathway activity and high intrinsic trametinib resistance.

**Figure 3.**
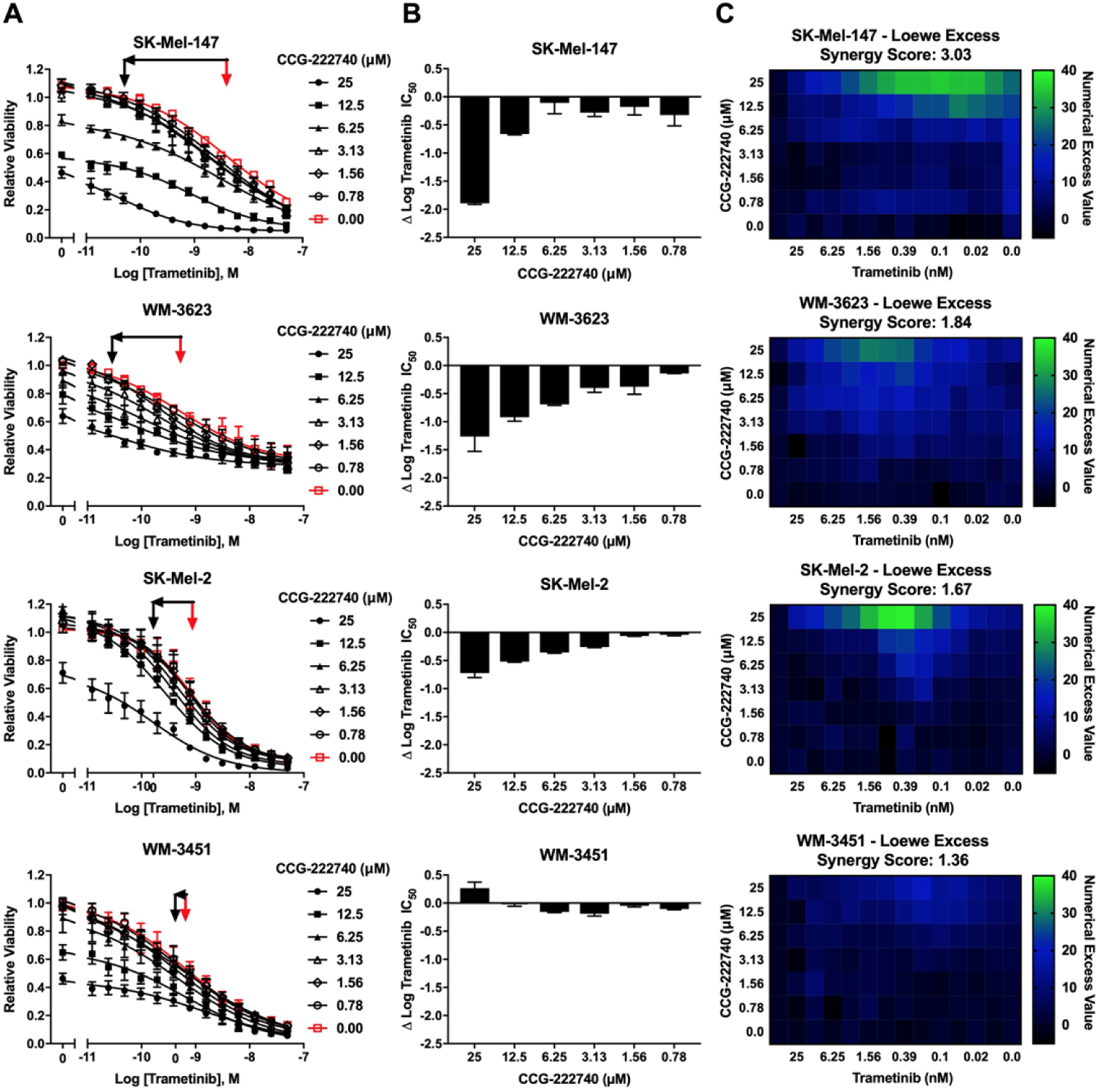
CCG-222740 synergizes with trametinib to inhibit NRAS mutant melanoma cell viability. **A.** Sensitivity to CCG-222740 and Trametinib in combination was determined by treating each cell line with increasing concentrations of trametinib in the presence of a range of concentrations of CCG-222740. Cell viability at 72 hours was determined using Cell-TiterGLO®. Values are expressed as the fraction of luminescence over vehicle control for at least three independent experiments. The red arrow indicates the logIC_50_ of trametinib in the absence of CCG-222740 and the black arrow indicates the logIC_50_ of co-treatment at 12.5 nM or 25 nM CCG-222740 which ever demonstrated the greatest leftward shift in the trametinib concentration response curve. **B.** ΔlogIC_50_ of Trametinib is determined as the difference between the logIC_50_ of trametinib in the absence and presence of CCG-222740. **C.** Loewe Excess was utilized as a metric of synergistic effects of the combination treatments. Loewe Excess was determined by detecting Numerical Excess Values using Chalice Analyzer. This software evaluates the observed data set compared to the predicted model of additivity to assess synergy. Expressed values represent the mean of at least three independent experiments.

### CCG-222740 disrupts nuclear localization of MRTF-A and MRTF-B but not YAP

As previously reported (20) for a structurally similar compound, CCG-203971, CCG-222740 effectively reduced MRTF-A nuclear localization in a concentration dependent manner in SK-Mel-147 cells (Fig. 4A-B). It also reduced the percentage of nuclear MRTF-B (Fig. 4C-D), however, there was no significant difference in the nuclear localization of YAP following CCG-222740 treatment (Fig. 4E-F). We also observed decreased mRNA levels of MRTF target genes CYR61, ANKRD1, CRIM1 and THBS1 upon 24-hour treatment of SK-Mel-147 cells with CCG-222740 in the RNA-Seq analysis (Supplementary Table S1), which was confirmed by qRT-PCR analysis (40-80% decrease, Supplementary Figure S3). These results are in accordance with inhibition of Rho/MRTF pathway.

**Figure 4.**
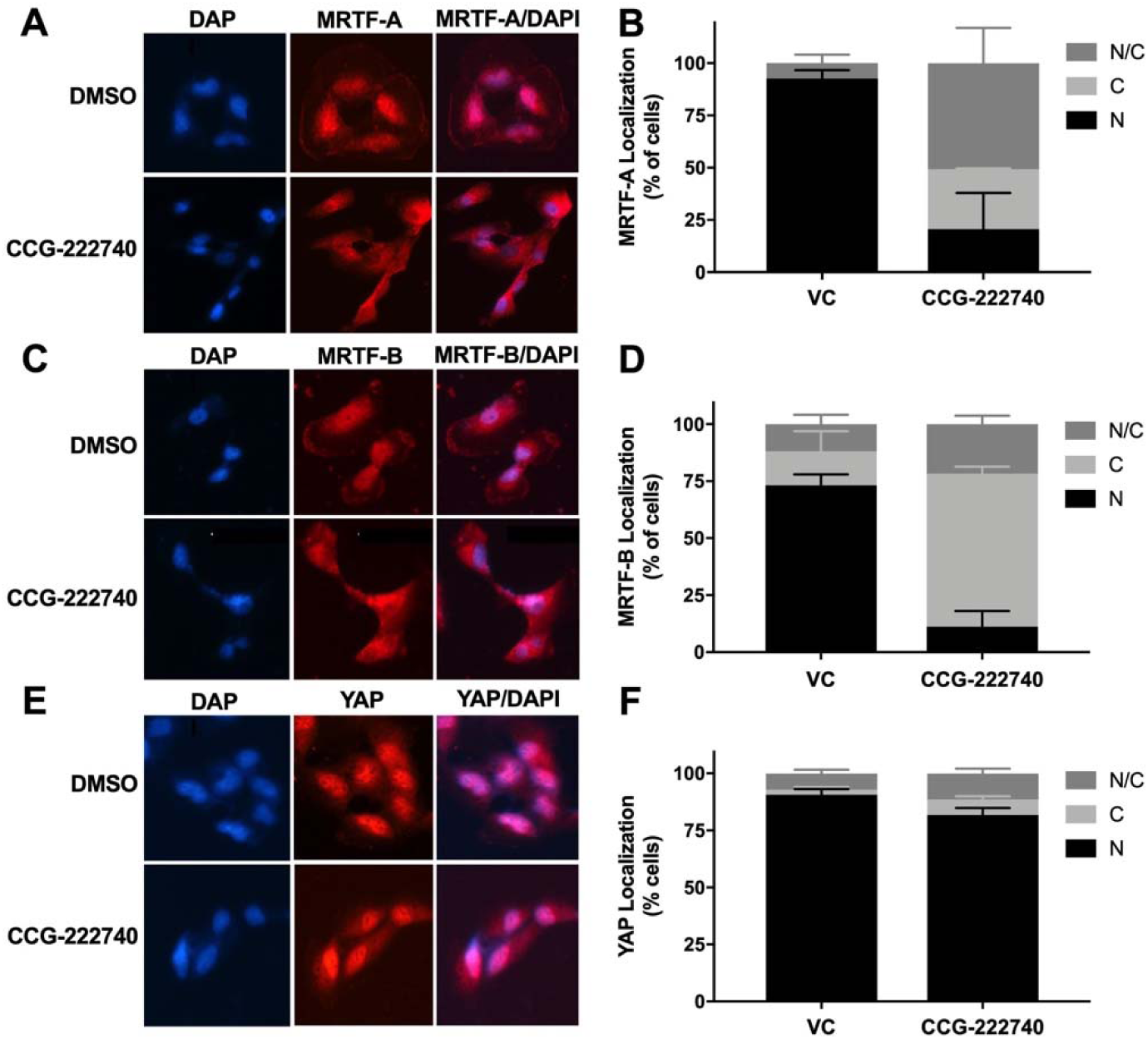
CCG-222740 disrupts nuclear localization of MRTF-A and MRTF-B but not YAP. **A.** Cellular localization of MRTF-A was determined in SK-Mel-147 cells using immunofluorescence following 24 hours treatment of vehicle control (VC) 0.1% DMSO or 10 µM CCG-222740 in the presence of 10% FBS. **B.** Quantification of MRTF-A cellular localization was determined by scoring individual cells as exclusively nuclear (N), cytosolic (C), or even distribution (N/C). Counts from two independent experiments with at least 100 cells scored blindly for each condition. **C.** Cellular localization of MRTF-B was determined in SK-Mel-147 cells as stated above for MRTF-A. **D.** Quantification of MRTF-B cellular localization was determined by scoring individual cells as exclusively nuclear (N), cytosolic (C), or even distribution (N/C). Counts from two independent experiments with at least 100 cells scored blindly for each condition. **E.** Cellular localization of YAP was determined in SK-Mel-147 cells as stated above for MRTF-A. **F.** Quantification of MRTF-B cellular localization was determined by scoring individual cells as exclusively nuclear (N), cytosolic (C), or even distribution (N/C). Counts from three independent experiments with at least 100 cells scored blindly for each condition.

### Potentiation of trametinib by CCG-222740 is specific to Rho/MRTF-pathway mechanisms

To determine whether potentiation of trametinib by CCG-222740 is specific to Rho/MRTF-pathway inhibition or merely a result of combination with a cytotoxic agent, we first tested trametinib in combination with vinblastine which is a microtubule assembly disrupting chemotherapeutic agent (39). Vinblastine alone, when used at 1-10 nM, efficiently decreased SK-Mel-147 cell viability, yet when used in combination with trametinib, there was no significant shift in the log IC_50_ of trametinib (Fig. 5A). Also, the synergy score was low, 1.42, for the combination of vinblastine and trametinib (Fig. 5B). As another agent to perturb Rho/MRTF signaling, we tested the effects of the ROCK inhibitor Y-27632 in combination with trametinib (40). ROCK signaling is upstream of MRTF activation and is a key component of the Rho/MRTF-pathway. When Y-27632 was used in combination with trametinib in SK-Mel-147 cells, there was a significant shift in log IC_50_, nearly one log difference compared to trametinib alone (Fig. 5C). The synergy score was also high at 2.11 based on Loewe Excess analysis (Fig. 5D). These results show that inhibition of the Rho/MRTF-pathway by two distinct agents is synergistic in combination with trametinib, while the effects of the non-targeted agent vinblastine and trametinib are additive.

**Figure 5.**
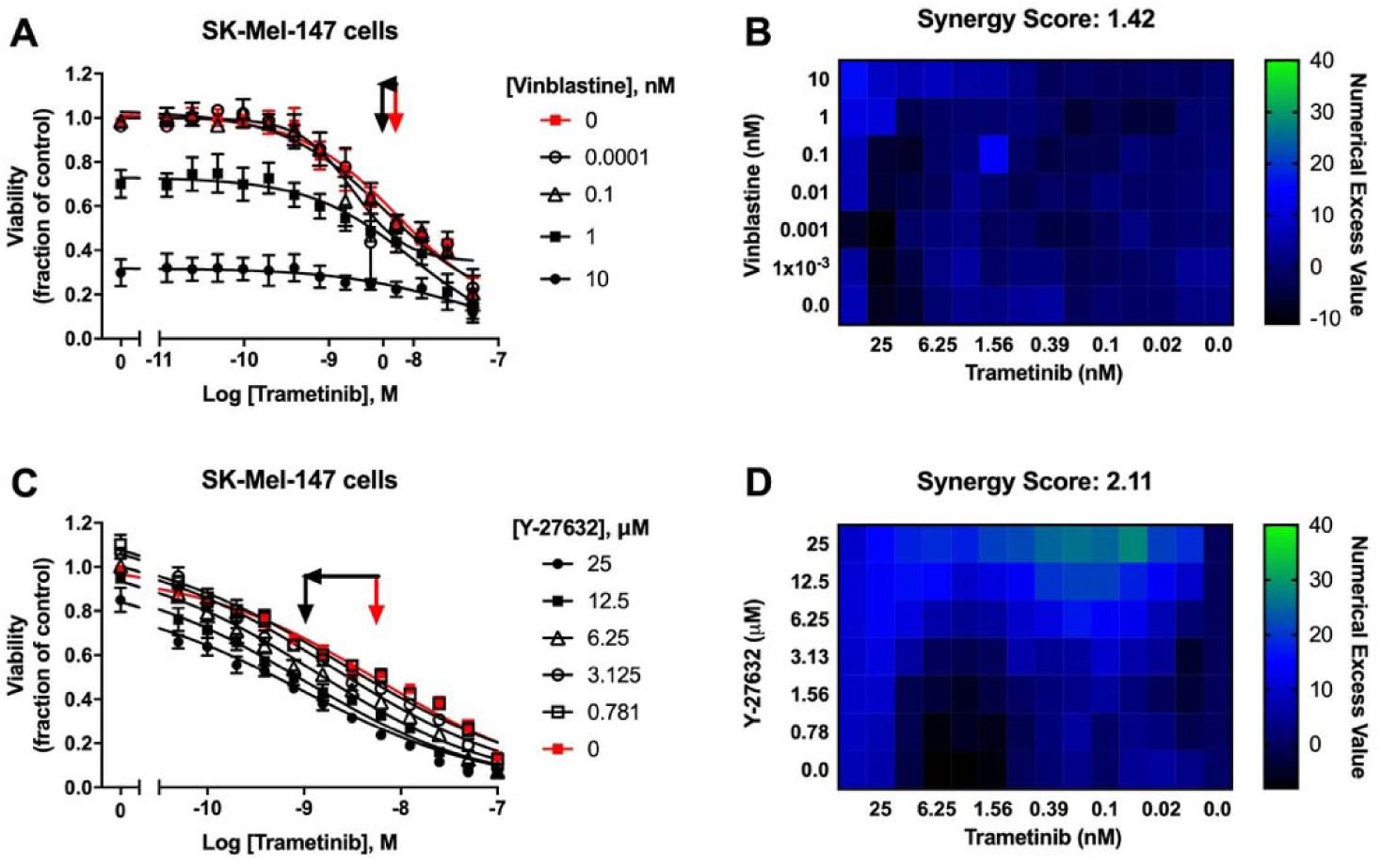
Potentiation of trametinib action is specific to the Rho/MRTF-pathway mechanisms. **A.** Sensitivity to trametinib in combination with Vinblastine. Combination treatments were studied in the same manner as described in Figure 3A. The reported values are expressed as the fraction of luminescence over vehicle control for three independent experiments. **B.** Loewe Excess was utilized as a metric of synergistic effects of the combination treatments in Figure 5A as described in Figure 3C. Expressed values represent the mean of three independent experiments. **C.** Sensitivity to trametinib in combination with the ROCK inhibitor Y-27632. Combination treatment was determined in the same manner as described above Figure 3A**. D.** Loewe Excess was determined for the combination treatments in Figure 5C as described in Figure 3C. Expressed values represent the mean of three independent experiments.

### Combination treatment with trametinib and CCG-222740 induces apoptosis and inhibits clonogenicity in SK-Mel-147 cells

To evaluate the functional effects of the combination treatment, we measured Caspase3/7 activity as a measure of apoptosis induction. CCG-222740 alone (10 µM) did not affect basal apoptosis as indicated by no change in caspase3/7 activity compared to control (Fig. 6A). Low nanomolar trametinib increased caspase3/7 activity compared to untreated cells, although the effect was not statistically significant (Fig. 6A). However, when trametinib was used in combination with CCG-222740, a significant increase in caspase3/7 activity was observed in SK-Mel-147 cells (Fig. 6A). These data provide evidence that SK-Mel-147 cells, which are highly resistant to trametinib-induced cell death, demonstrate cooperative induction of apoptosis when the Rho/MRTF-pathway inhibitor CCG-222740 is used in combination.

**Figure 6.**
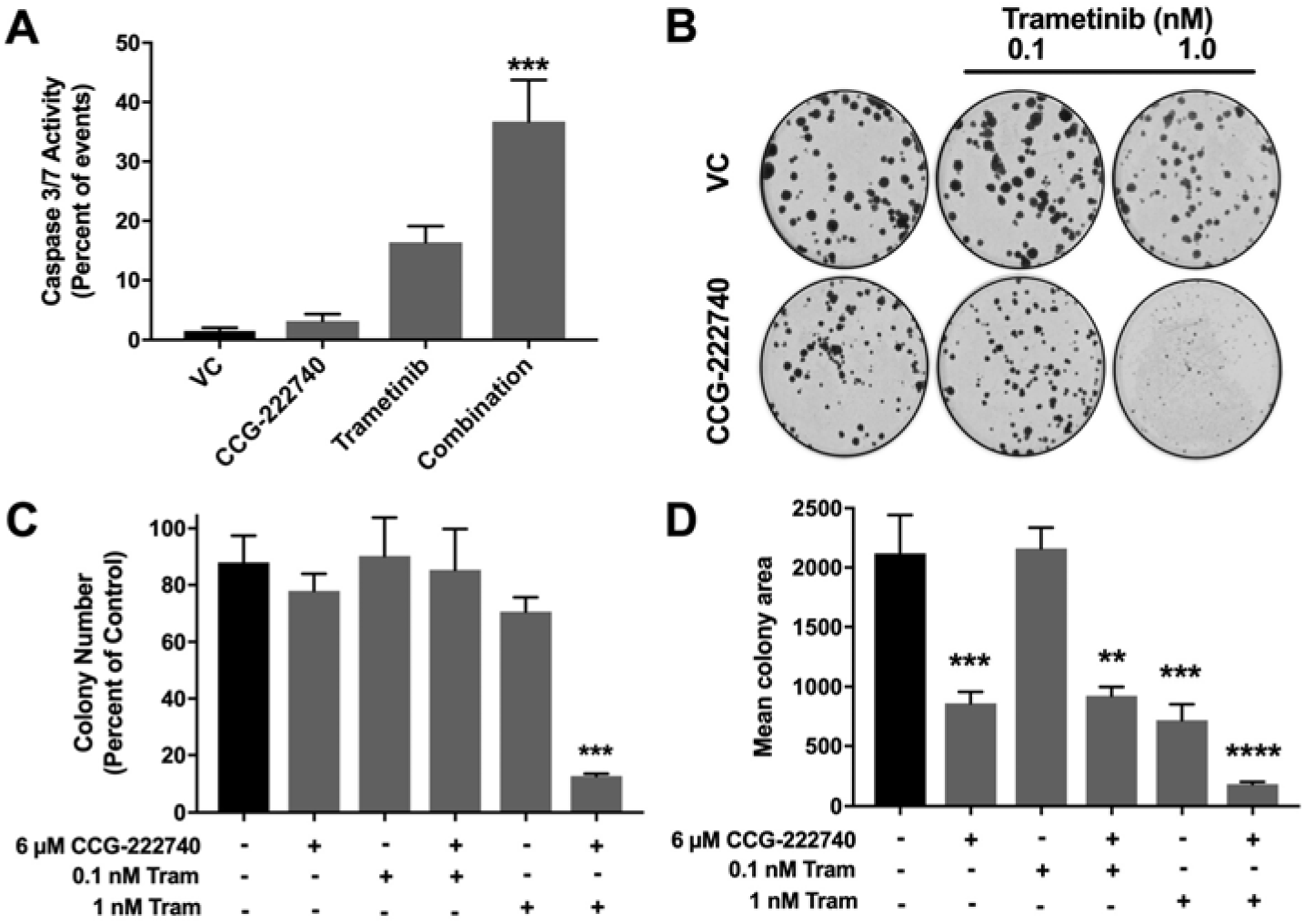
Combination treatment with trametinib and CCG-222740 induces apoptosis and inhibits clonogenicity in SK-Mel-147 cells. **A.** Caspase 3/7 activity was determined using flow cytometry 48 hours after treatment with 10 µM CCG-222740 and 12.5 nM Trametinib alone or in combination and 0.2% DMSO VC. **B.** SK-Mel-147 cells were seeded at low density (200 cells/well) into 6-well plates and simultaneously treated with DMSO control, 6 µM CCG-222740, 0.1 nM Trametinib, 1 nM Trametinib, or a combination of each. Shown are the representative images of colonies formed under indicated treatment conditions. **C.** Number of colonies were counted using ImageJ with a cutoff for colony size (pixel^2) fixed at 50-infinity and circularity defined as 0.2-1.0. Results are expressed as the mean (±SEM) of triplicate experiments (***p<0.005 vs. VC). **D.** Mean colony area was determined using ImageJ and the results are expressed as the mean (±SEM) of triplicate experiments (**p<0.01, ***p<0.005, ****p<0.0001 vs. VC).

Finally, we performed a colony formation assay to assess clonogenicity of SK-Mel-147 cells in the presence of trametinib and CCG-222740. Treatment with CCG-222740 alone (6 µM) or trametinib alone (1 nM) had no significant effect on the number of colonies formed by SK-Mel-147 cells (Fig. 6B). However, when cells were treated with 1 nM trametinib and 6 µM CCG-222740 in combination, colony formation was nearly eliminated (Fig. 6B-C). When we evaluated mean colony area, we detected significant decreases across single agent and combination treatments except for 0.1 nM trametinib alone (Fig. 6D). The greatest decrease in mean colony area was detected when cells were treated with 1 nM trametinib in combination with 6 µM CCG-222740 (Fig 6D). Thus, combination treatment with trametinib and CCG-222470 markedly inhibited clonogenicity of SK-Mel-147 cells.

## DISCUSSION

Given the current lack of approved targeted therapies for NRAS mutant melanoma patients, MEK inhibitors offer a potential treatment option. However, intrinsic and acquired resistance to these inhibitors is a major hurdle. In a recent study, Najem et al. observed that MEK inhibition using pimasertib had only a limited effect on apoptosis in NRAS^Q61^ mutant melanoma cell lines (14). They explained this by a systematic MITF upregulation and resultant upregulation of Bcl-2 upon pimasertib treatment. Here, we show that a panel of four NRAS^Q61^ mutant melanoma cell lines had varying sensitivity to trametinib treatment and that the Rho/MRTF pathway was activated in the subset of NRAS mutant melanoma cell lines with high intrinsic resistance to trametinib-mediated cell growth inhibition.

Amplification of RhoA/C or MKL1/2 (gene names for MRTF-A/B) or mutations in upstream activators of RhoA/C have been found in ∼30% cutaneous melanomas. Increased expression of RhoC or MRTF-A has been linked to aggressive disease and overall poor patient survival (20). While the MRTF-SRF transcriptional axis plays a pro-metastatic role in cancers, the Rho/MRTF pathway is also emerging as a drug resistance mechanism in different types of skin malignancies. Whitson et al. reported that activation of the Rho/MRTF pathway promoted resistance to a smoothened (SMO) inhibitor in basal cell carcinoma (BCC) (41). Treatment with the MRTF pathway inhibitors CCG-1432 and CCG-203971 had considerable efficacy in treating resistant BCC *in vivo* indicating a therapeutic potential of this pathway in drug-resistant malignancies. Additionally, we recently reported Rho/MRTF pathway activation in a subset of BRAF inhibitor-resistant BRAF^V600^ mutant melanoma cell lines (34). Inhibition of this pathway via CCG-222740 re-sensitized the resistant cells to vemurafenib. Here, we observed greater activation of the Rho/MRTF pathway in SK-Mel-147 and other NRAS mutant melanoma cells compared to the BRAF mutant SK-Mel-19 cells, as measured by stress fiber formation, nuclear localization of MRTF-A/B and mRNA expression levels of a target gene, CYR61. Importantly, Rho/MRTF pathway activation correlated strongly with intrinsic resistance to trametinib in these NRAS mutant cell lines. In light of the inverse relationship between pERK and Rho/MRTF activation, it is plausible that the Rho/MRTF pathway takes over for the MAPK pathway in driving cell proliferation and survival.

Considering the lack of success of therapies targeting BRAF and MEK in NRAS mutant melanoma patients (42), recent efforts have been focused on finding other therapeutic targets and development of combination therapies. For example, inhibiting a novel NRAS-activating kinase (STK19) and combining MEK inhibitors with inhibitors of the MER receptor tyrosine kinase (MERTK), BET, and HDAC have been reported to block NRAS mutant melanoma growth *in vitro* and *in vivo* (15-17,43). In this study, we demonstrate that the combination of a Rho/MRTF pathway inhibitor, CCG-222740 and the MEK inhibitor trametinib synergistically inhibited viability of a subset of NRAS mutant melanoma cells and also induced apoptosis in cell with highly activated Rho/MRTF pathway. Similar results on cell viability were obtained when trametinib was used in combination with an inhibitor of ROCK, which is an important component of the Rho/MRTF pathway. Our results are in accordance with previous reports which demonstrated that a combination of MEK and ROCK inhibitors not only reduced NRAS mutant melanoma cell viability *in vitro* but also reduced tumor growth *in vivo* (12). In SK-MEL-147 cells, which are the most trametinib-resistant cells in our panel, 12.5 nM trametinib in combination of 10 µM CCG-222740 was required to induce apoptosis. This trametinib dose is consistent with or lower than other *in vitro* studies involving synergistic combination of trametinib with another small molecule inhibitor (14,16,44). For example, Vogel et al. reported cooperative induction of apoptosis in SK-MEL-147 cells when a combination of 100 nM trametinib and 1 µM ROCK inhibitor GSK269962A was used (12). Remarkably, only 1 nM trametinib was needed along with 6 µM CCG-222740 to dramatically suppress clonogenicity of the SK-Mel-147 cells. Also, 12.5 nM trametinib and 10 µM CCG-222740 induced apoptosis in SK-Mel-147 cells. Use of such low MEK inhibitor concentrations is a significant benefit, because severe dose-limiting toxicity is a concern with MEK inhibitors.

We observed that CCG-222740 disrupted nuclear localization of MRTF in SK-Mel-147 cells, which was accompanied by downregulation of MRTF target genes CYR61, ANKRD1, CRIM1 and THBS1. These genes in the MRTF-SRF axis could also be regulated by YAP, a transcription factor in the Hippo pathway. Using CYR61 as a model target gene, Yu et al. demonstrated that activation of both MRTF-A and YAP pathways and functional interactions between MRTF-A and YAP are required for transcriptional control of RhoA-regulated genes (45). Foster et al. further showed that expression of MRTF-SRF target genes and expression of YAP target genes are interdependent, even when only one of these pathways directly regulates the target gene (46). They further found that activation of one of these pathways indirectly activates the other pathway depending on the cytoskeletal dynamics. Here, we found that YAP was localized in the nucleus of all the cell lines tested, irrespective of the BRAF or NRAS mutational status and sensitivity of cells to trametinib. Additionally, YAP remained localized in the nucleus even after the treatment with CCG-222740. We attempted to investigate the involvement of YAP and Hippo pathway in potentiation of trametinib by CCG-222740 by using verteporfin, an inhibitor of YAP (47). However, we were unable to obtain consistent and reproducible results in the cell viability assay when testing the effects of combination of verteporfin and trametinib (data not shown).

In conclusion, we report a role of the Rho/MRTF pathway in intrinsic resistance of NRAS mutant melanoma cells to MEK inhibitor-induced cell death. CCG-222740, a Rho/MRTF pathway inhibitor, markedly increased the activity of trametinib on NRAS-mutant melanoma cell lines. The combination of CCG-222740 and trametinib induced apoptosis and inhibited clonogenicity in those NRAS mutant melanoma cell lines having increased Rho/MRTF activation. These results warrant further *in vivo* studies and potentially clinical development of combinations of MEK inhibitors with Rho/MRTF pathway inhibitors.

## Conflict of interest

Dr. Neubig is Founder and President of FibosIX, Inc. which has a license option for CCG-222740.

## Acknowledgements

We would like to thank Dr. Erika Lisabeth, Dr. William Jackson, Dr. Cheryl Rockwell and Dr. Kathleen Gallo for helpful discussions.

## Notes

Financial support: MSU Gran Fondo Skin Cancer Research Fund, NIH F31 CA232555 (SAM)

