## Supplemental_Information for "Inhibition of the Myocardin-Related Transcription Factor pathway increases efficacy of Trametinib in NRAS-mutant melanoma cell lines"

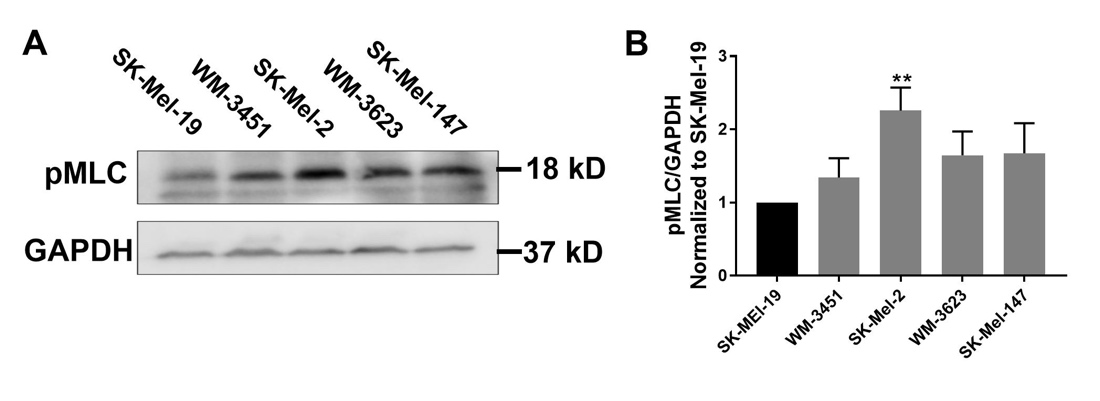

**Supplementary Figure S1**. **A.** Immunoblots were conducted across the melanoma cell line panel to detect pMLC or GAPDH as a protein loading control. **B.** Quantitative band density analysis was performed for each experiment comparing the intensity of pMLC relative to control GAPDH. Results are expressed as the mean (±SEM) of triplicate experiments (**p<0.01 vs. SK-Mel-19).

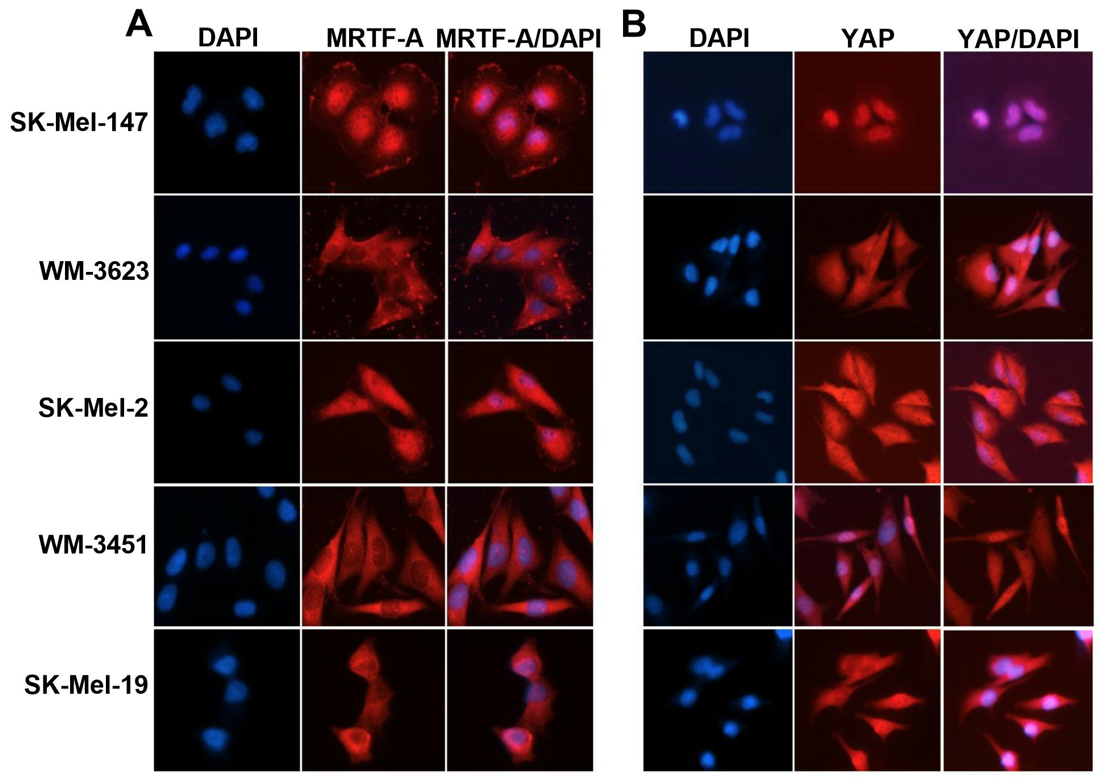

**Supplementary Figure S2**. **A.** Representative images of the cellular localization of MRTF-A using immunofluorescence in melanoma cell lines (10% FBS). **B.** Representative images of the cellular localization of YAP using immunofluorescence in melanoma cell lines (10% FBS).

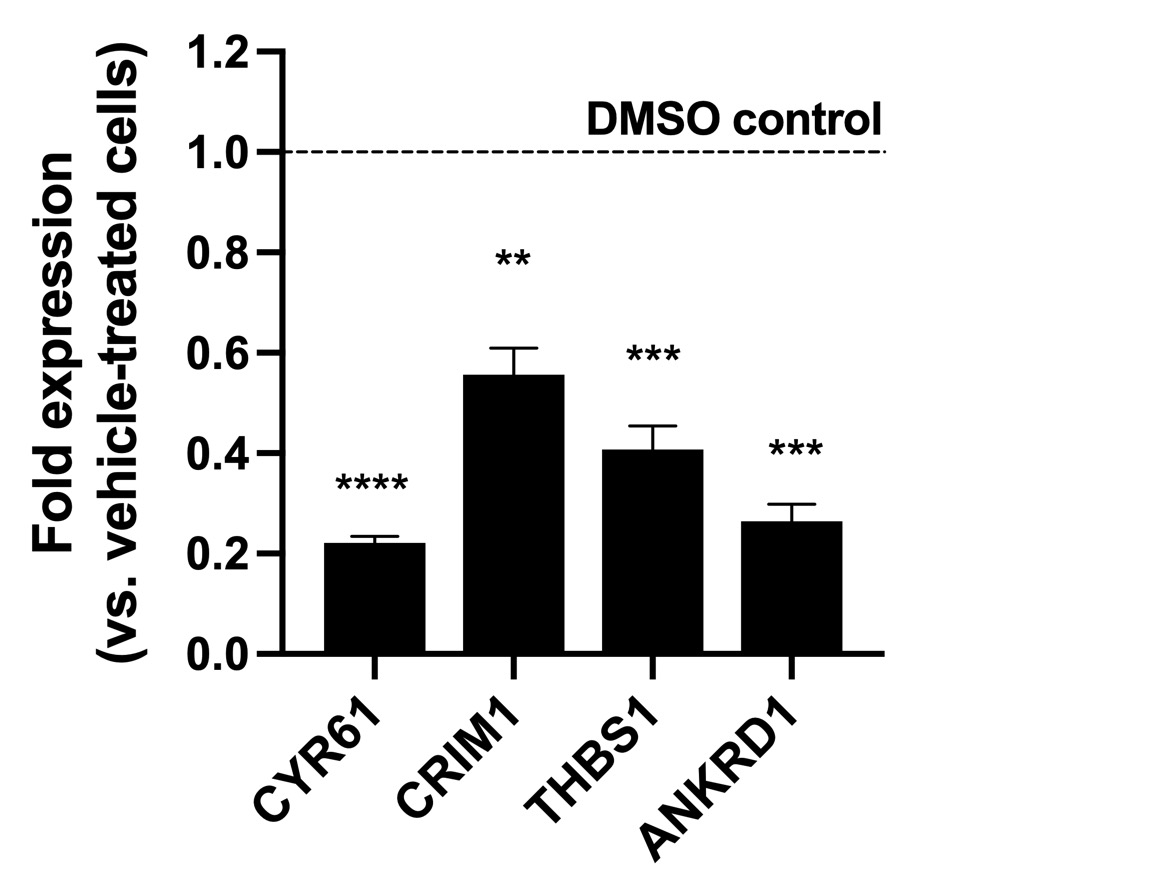

**Supplementary Figure S3**. CCG-2222740 treatment reduces expression of MRTF target genes CYR61, CRIM1, THBS1 and ANKRD1. SK-Mel-147 cells were treated for 24 hours with 10 µM CCG-222740 or DMSO (0.02%) before isolation of RNA for qPCR analysis. Expression of target genes were normalized against GAPDAH and results are expressed as the mean ( ±SEM) of triplicate experiments (*, P < 0.01, **, P < 0.001, ***, P < 0.0001, ****, P < 0.00001 vs. DMSO control).

| **Ensembl ID** | **Approved Symbol** | **logFC** | **logCPM** | **PValue** | **FDR** |
| --- | --- | --- | --- | --- | --- |
| **ENSG00000168003** | SLC3A2 | 1.740277527 | 8.507639444 | 1.21E-14 | 7.71E-10 |
| **ENSG00000115380** | EFEMP1 | -1.706565216 | 3.530197807 | 9.55E-11 | 3.04E-06 |
| **ENSG00000091986** | CCDC80 | -1.495390124 | 4.347019002 | 4.67E-09 | 9.92E-05 |
| **ENSG00000181449** | SOX2 | 2.140598107 | 0.239972409 | 3.44E-08 | 0.000547402 |
| **ENSG00000104419** | NDRG1 | 1.363870834 | 5.978803232 | 4.66E-08 | 0.000586459 |
| **ENSG00000148677** | ANKRD1 | -1.397055415 | 4.627081746 | 5.59E-08 | 0.000586459 |
| **ENSG00000150938** | CRIM1 | -1.308149346 | 6.808500344 | 6.45E-08 | 0.000586459 |
| **ENSG00000224773** | HSPA8P7 | -1.98036841 | 2.305467038 | 1.86E-07 | 0.001477206 |
| **ENSG00000137801** | THBS1 | -1.185139541 | 10.08725598 | 2.85E-07 | 0.00201518 |
| **ENSG00000196878** | LAMB3 | 1.137023364 | 7.353579658 | 9.04E-07 | 0.005469966 |
| **ENSG00000213820** | RPL13P2 | -2.378109738 | 2.097258073 | 9.45E-07 | 0.005469966 |
| **ENSG00000128564** | VGF | 1.413051246 | 1.555022873 | 1.3E-06 | 0.006602879 |
| **ENSG00000154734** | ADAMTS1 | -1.102716883 | 6.357359483 | 1.35E-06 | 0.006602879 |
| **ENSG00000236105** | PRELID3BP10 | -2.521855225 | -0.667607432 | 1.89E-06 | 0.008593398 |
| **ENSG00000231445** | TIMM8AP1 | -2.03756353 | -0.277981649 | 2.78E-06 | 0.01016252 |
| **ENSG00000129757** | CDKN1C | 1.19260645 | 3.011058803 | 2.85E-06 | 0.01016252 |
| **ENSG00000142871** | CYR61 | -1.036485735 | 9.347827715 | 2.87E-06 | 0.01016252 |
| **ENSG00000092969** | TGFB2 | -1.146679925 | 4.714269799 | 3.1E-06 | 0.010396226 |
| **ENSG00000052841** | TTC17 | 1.075378747 | 6.169282608 | 5.58E-06 | 0.017771644 |
| **ENSG00000078401** | EDN1 | -1.202295537 | 3.121136369 | 5.94E-06 | 0.018013191 |
| **ENSG00000052802** | MSMO1 | 1.079766996 | 5.379290384 | 7.15E-06 | 0.020689773 |
| **ENSG00000138798** | EGF | -1.282835697 | 2.550065363 | 7.79E-06 | 0.021564115 |
| **ENSG00000170439** | METTL7B | 1.152224231 | 2.41811322 | 1.06E-05 | 0.028090638 |
| **ENSG00000114251** | WNT5A | -1.050718673 | 5.271773994 | 1.39E-05 | 0.032740269 |
| **ENSG00000232480** | TGFB2-AS1 | -1.851166895 | 0.032554217 | 1.4E-05 | 0.032740269 |
| **ENSG00000168028** | RPSA | -1.157332873 | 6.684989777 | 1.44E-05 | 0.032740269 |
| **ENSG00000145632** | PLK2 | -0.9729373 | 7.098792127 | 1.44E-05 | 0.032740269 |
| **ENSG00000151892** | GFRA1 | -1.241962153 | 1.757886426 | 1.67E-05 | 0.035350063 |
| **ENSG00000170523** | KRT83 | -1.706326252 | 0.010133492 | 2.25E-05 | 0.045127787 |
| **ENSG00000214274** | ANG | 1.837466378 | -0.700807455 | 2.27E-05 | 0.045127787 |

**Supplementary Table S1**. RNA-Seq analysis identifies differential gene expression in SK-Mel-147 cells upon 24-hour treatment with CCG-222740, compared to untreated SK-Mel-147 cells. Negative logFC values indicate genes downregulated upon CCG-222740 treatment, and positive values indicate genes upregulated upon CCG-222740 treatment.
